# DeepDRIM: a deep neural network to reconstruct cell-type-specific gene regulatory network using single-cell RNA-Seq Data

**DOI:** 10.1101/2021.02.03.429484

**Authors:** Jiaxing Chen, Chinwang Cheong, Liang Lan, Xin Zhou, Jiming Liu, Aiping Lyu, William K Cheung, Lu Zhang

## Abstract

Single-cell RNA sequencing is used to capture cell-specific gene expression, thus allowing reconstruction of gene regulatory networks. The existing algorithms struggle to deal with dropouts and cellular heterogeneity, and commonly require pseudotime-ordered cells. Here, we describe DeepDRIM a supervised deep neural network that represents gene pair joint expression as images and considers the neighborhood context to eliminate the transitive interactions. DeepDRIM yields significantly better performance than the other nine algorithms used on the eight cell lines tested, and can be used to successfully discriminate key functional modules between patients with mild and severe symptoms of coronavirus disease 2019 (COVID-19).

## Background

Reconstruction of gene regulatory networks (GRNs) is critical to understand the mechanisms of synergic gene effects and context-specific transcriptional dynamics. High-throughput technologies such as chromatin immunoprecipitation (ChIP)-chip and ChIP-seq can directly capture the transcription factor (TF) binding sites of targeted genes; however, these techniques are costly and TF-specific, and are therefore unsuitable for use on a whole-genome scale [45]. As a consequential observation, the fact that the co-expression of TFs and their target genes has been adopted to reconstruct GRNs [22, 17, 32, 26] as a means of reverse engineering. In the last two decades, microarrays and bulk RNA sequencing (RNA-seq) have been the two mainstream technologies used to capture gene expression profiles from diverse tissues. Both techniques have been widely applied to identify differentially expressed genes and reconstruct GRNs [43, 25]. However, microarrays and RNA-seq inappropriately assume that gene expression is homogeneous among cells and ignore cellular heterogeneity. Indeed, tissue consists of a diverse range of cell types with distinct GRNs [20] and biological functions [7]. Several studies have sought to reconstruct GRNs using bulk gene expression data [24, 53], but the cell-type-specific GRNs remain largely unexplored. Single-cell RNA sequencing (scRNA-seq) offers an opportunity to capture cell-specific gene expression, which in turn could provide deeper insights into the cellular heterogeneity and cell-type-specific gene activities[51].

Most of the available algorithms for GRN reconstruction are designed for bulk gene expression, and function by resolving two computational challenges. In this context, unique difficulties arise if scRNA-seq data are adopted instead. First, putative TF-gene interactions are derived by examining their co-expression. Bulk gene expression data are commonly normalized to a standard Gaussian distribution, such that the TF-gene correlation can be quantified by methods such as mutual information (MI) [37], Pearson correlation coefficient (PCC) [48, 40]. The scRNA-seq gene expression data are zero-inflated due to the imbalanced transcript sampling. Although it is possible to impute zero entries before calculating the TF-gene co-expression, this may introduce unpredictable noise and bias [4], given that most of the imputation algorithms make use of gene-gene co-expression. Second, the TF-gene pairs with strong co-expression due to transitive interactions (e.g., those bridged by one or more intermediate genes) should be eliminated (Additional file 1: Figure S1). Several strategies have been designed to remove these transitive interactions by conditioning on the other confounding genes; examples include the Gaussian graphical model [3], conditional MI [62], context-based normalization and edge removal [17], and tree-based ensemble methods [26]. Unfortunately, these algorithms were originally developed to analyze bulk gene expression data, and are unsuitable for modeling scRNA-seq data [10]. Many algorithms have recently been proposed to cater for the unique characteristics of scRNA-seq for GRN reconstruction. SCODE [38] infers cell-specific pseudo-time and reconstructs the GRN by solving ordinary differential equations. PIDC [9] adopts partial information decomposition to break down the TF-gene correlation into redundant, synergistic, and unique effects. SINCERITIES [44] utilizes regularized linear regression to infer GRNs from time-stamped scRNA-seq data by referring to temporal changes in the gene expression distributions. GENIE3 [26] is a tree-based ensemble method that was initially developed for bulk gene expression data. Aibar *et al.* later applied GENIE3 to reconstruct the global GRN for scRNA-seq and developed AUCell to score the active gene signatures for each cell [1]. Although these dedicated strategies have been designed to deal with the inherent issues in scRNA-seq data, none of them yield acceptable results benchmarked by cell-type-specific ChIP-seq data, and some are even close to random guessing [46].

CNNC [60] is a supervised deep neural network that represents the joint expression of a gene pair as an image and uses convolutional neural networks (CNNs) to predict gene-gene co-expression from scRNA-seq data. CNNC is robust to dropouts and can infer the interaction causalities using the information from cell-type-specific ChIP-seq data. We generated synthetic GRNs and their corresponding gene expression data (**Methods** and Fig.1) to examine whether CNNC could effectively distinguish direct and transitive interactions. We noted that a substantial number of the false positives obtained with CNNC were centered in the gene pairs with strong Pearson correlations (Fig.1 A).

**FIG. 1.**
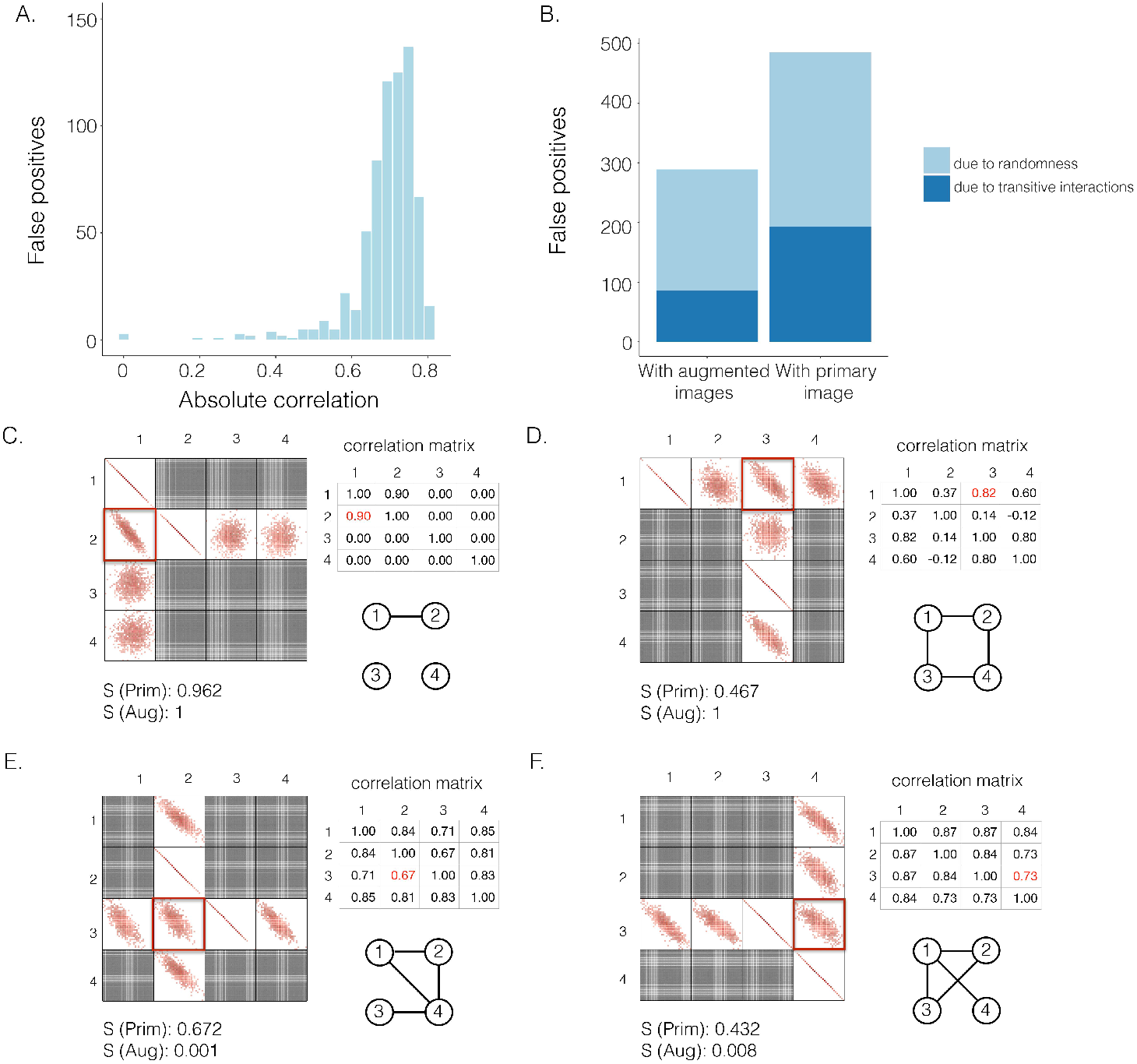
The effectiveness of neighbor images in reconstructing GRN on the simulated data. **A**. The distribution of false positives from CNNC. **B**. The false positives of the two models with primary (*Prim*) and augmented (*Aug*) images as inputs due to randomness and transitive interactions. **C** and **D**. Two examples that demonstrate both of the models can correctly identify the direct interactions (**C**: *g*_1_ ⇒ *g*_2_, **D**: *g*_1_ ⇒ *g*_3_). **E** and **F**. Two examples that demonstrate the model trained by augmented images can recognize and eliminate the false positives caused by the transitive edges (**E**: *g*_2_ ⇒ *g*_3_, **F**: *g*_3_ ⇒ *g*_4_). The primary images are highlighted in the red squares, and *S*(·) denotes the confidence scores from the models with primary (*S*(*Prim*)) or augmented images (*S*(*Aug*)) as input. The correlation matrices and absolute correlation are calculated by Pearson correlation coefficient.

Yet considering the image of the target TF-gene pair (primary image) as the only input for the prediction is insufficient (Fig.1 A). Inspired by an approach named context likelihood of relatedness (CLR) [17] which have been used to remove the transitive interactions by normalizing the MI of the target TF-gene pairs to z-scores with their corresponding neighborhood, one can in fact consider both the target TF-gene pair (primary image) and the images from the gene pairs that share one gene with the target pair (neighbor images) as the input to the model (Fig.1 and 2).

**FIG. 2.**
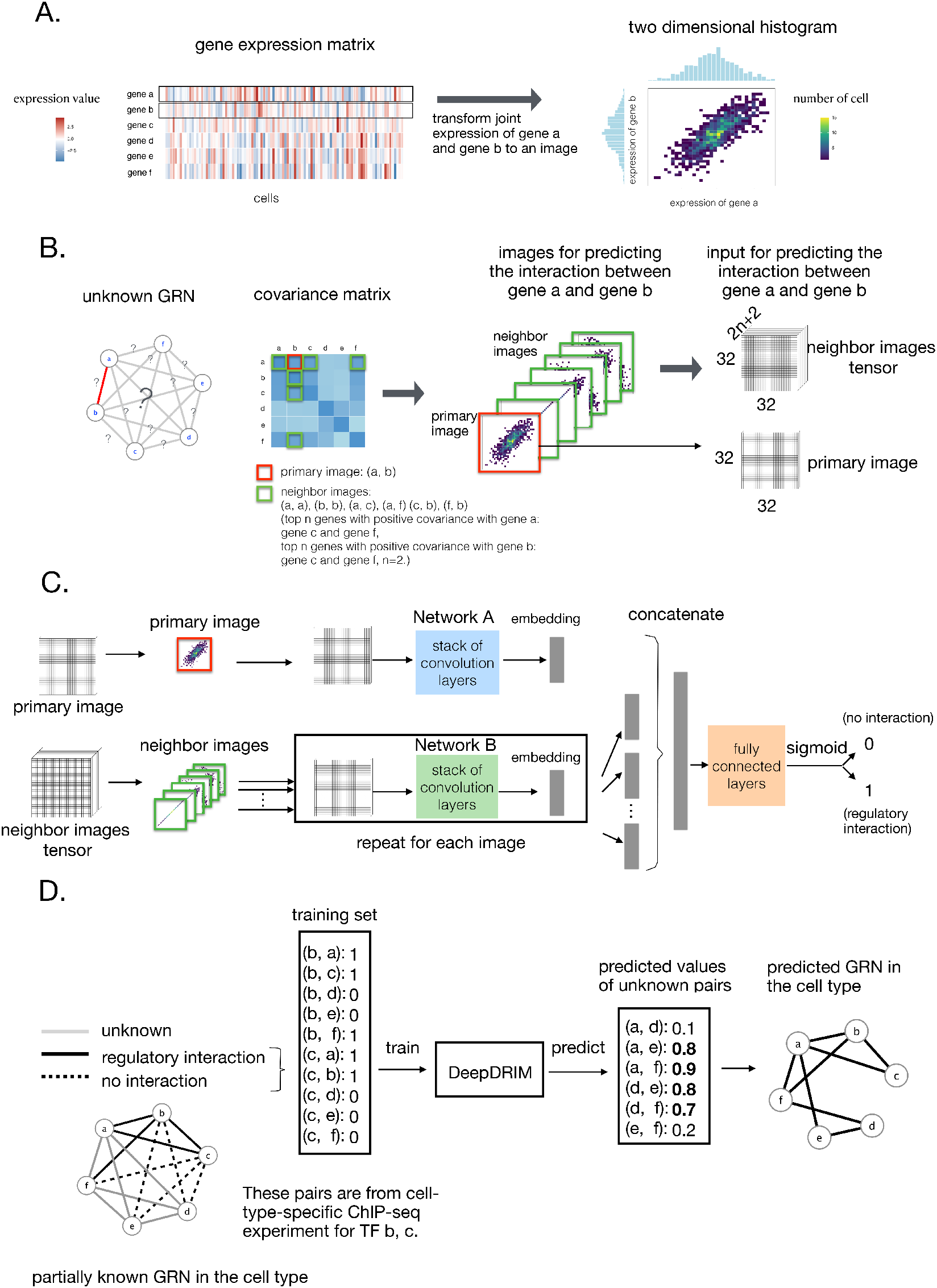
Overview of DeepDRIM. **A**. Representation of the joint gene expression of gene a and gene b as a primary image. **B**. The 2*n* + 2 neighbor images are generated from the genes with strong positive covariance with gene a or gene b. **C**. The network architecture of DeepDRIM, including Network A and Network B, which are two stacked convolutional embedding structures designed to process the primary and neighbor images, respectively. Detailed network structures are shown in Additional file 1: Figure S3. **D**. An example for the prediction of a cell-type-specific GRN using DeepDRIM.

Here we propose DeepDRIM (deep learning-based direct regulatory interaction model), a supervised deep neural network that can reconstruct highly accurate cell-type-specific GRNs from scRNA-seq data by considering both primary and neighbor images. The rationale and workflow of DeepDRIM are shown in Fig.2. DeepDRIM first transforms the primary and neighbor images (Fig.2 A and B) into low-dimensional embeddings using multiple convolutional layers, where their embeddings are then concatenated as the input to a multiple-layer perceptron to calculate the regulatory confidence scores (Fig.2 C). We compared the effectiveness of DeepDRIM with PCC, MI, GENIE3, and CNNC for the analysis of eight real scRNA-seq datasets. Our results demonstrated that DeepDRIM yielded the best performance with respect to both the area under the receiver operating characteristic curve (AUROC) and the area under the precision-recall curve (AUPRC), and significantly outperformed CNNC (Fig.3 A-D). We also compared DeepDRIM with six effective algorithms that were recently highlighted for reconstructing GRN on scRNA-seq data [46]. The results demonstrated that DeepDRIM substantially outperformed these algorithms on the five scRNA-seq datasets with the pseudotime-ordered cells (Fig.3 E-F). Further simulation demonstrated that the performance of DeepDRIM could be improved by involving more neighbor images, and was robust to the dropout rate, the cell number, and the size of the training set (Fig.4 A-D).

**FIG. 3.**
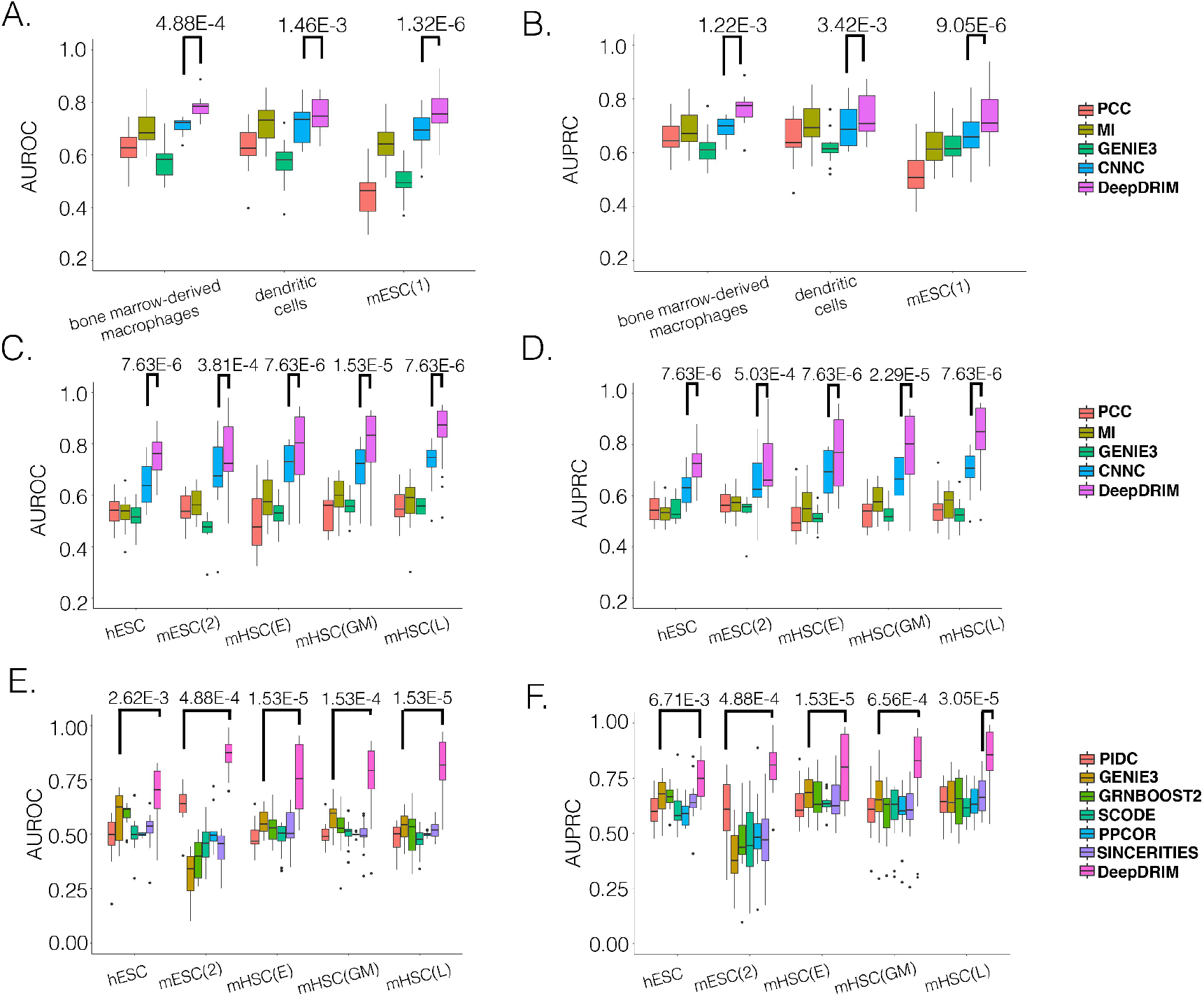
Comparison of DeepDRIM with the existing algorithms for GRN reconstruction on the scRNA-seq data from eight cell lines. **A**, **B**, **C** and **D**: *p*-values were calculated between CNNC and DeepDRIM. **E** and **F**: The *p*-values were calculated between DeepDRIM (the best performer) and the second best algorithms (Additional file 1: Table S3 and S4).

**FIG. 4.**
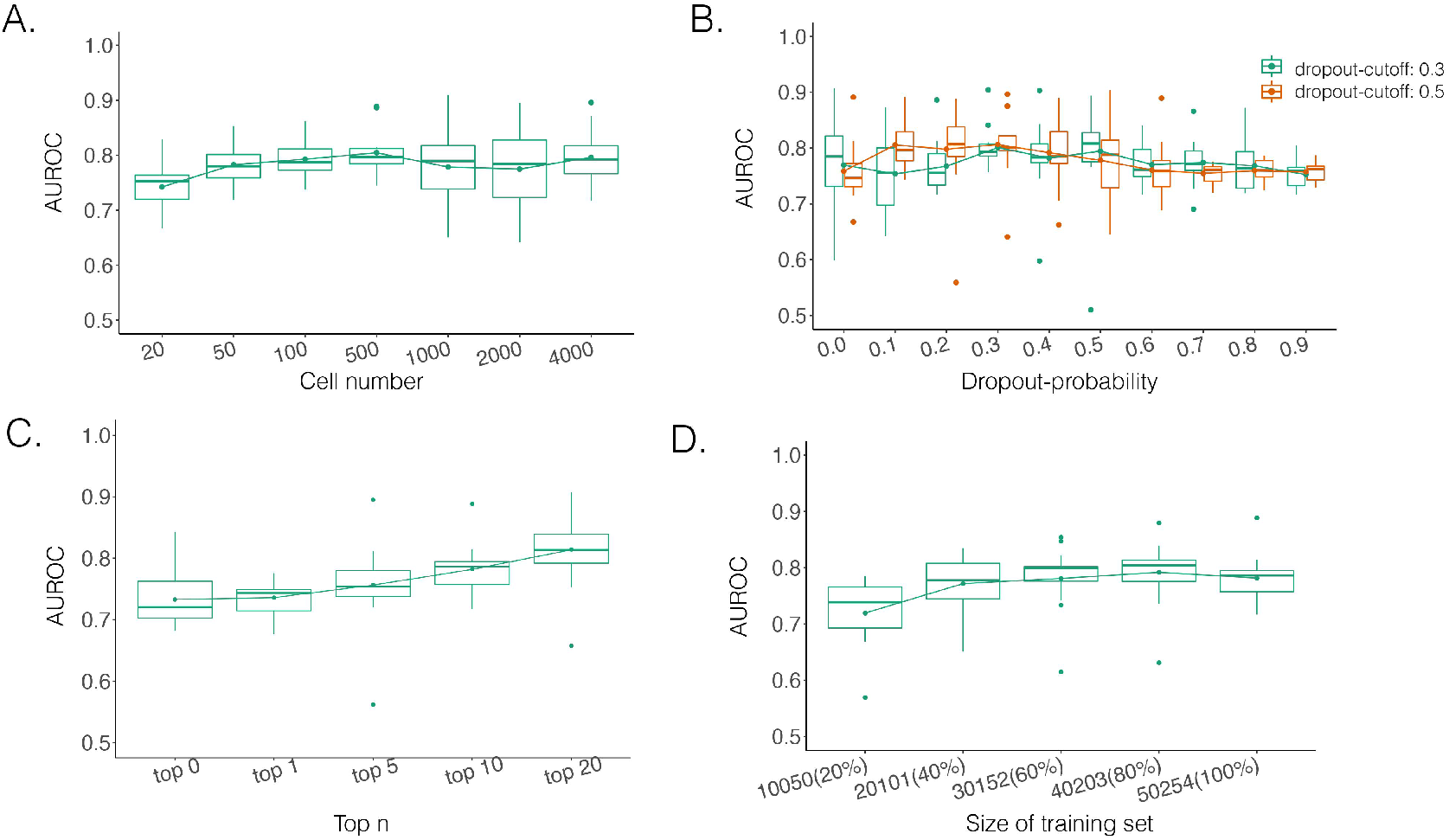
Performance of DeepDRIM with a wide range of the qualities of scRNA-seq data (cell numbers and dropout rates), the number of involved neighbor images, and the size of training set.

We applied DeepDRIM to the scRNA-seq data collected from the bronchoalveolar lavage fluid of patients with mild and severe symptoms of coronavirus disease 2019 (COVID-19) [34] to discover the changes in B cell-specific GRNs. As a result, we observed that a large number of differentially expressed TFs (DETFs) were “activated” in patients with severe disease (Fig.5 A and B). Furthermore, in patients with severe COVID-19 symptoms, the functions of the target genes were enriched in apoptosis, response to decreased oxygen levels, and microtubules (Fig.5 C, Fig.6 A and B), all of which have been previously shown to be associated with COVID-19 [8, 57] and virus infection [18].

**FIG. 5.**
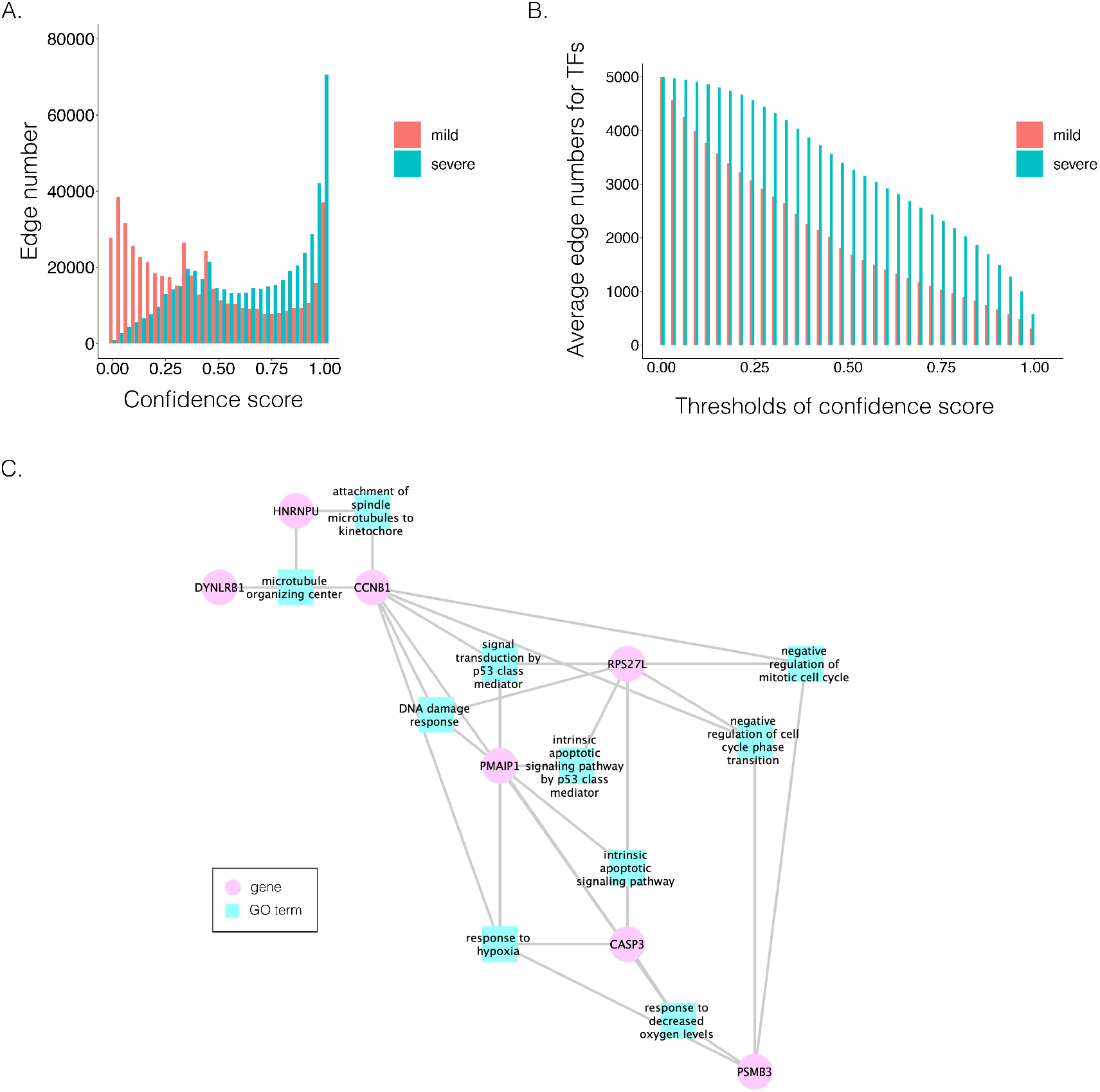
Comparison of B cell-specific gene regulatory networks (GRNs) for patients with mild and severe COVID-19. **A**. The distribution of the confidence scores of the differentially expressed transcription factors and their target genes. **B**. The average target numbers of DETFs given different confidence score thresholds. **C**. The GO modules and the involved key transcription factors/genes related to COVID-19 symptoms.

**FIG. 6.**
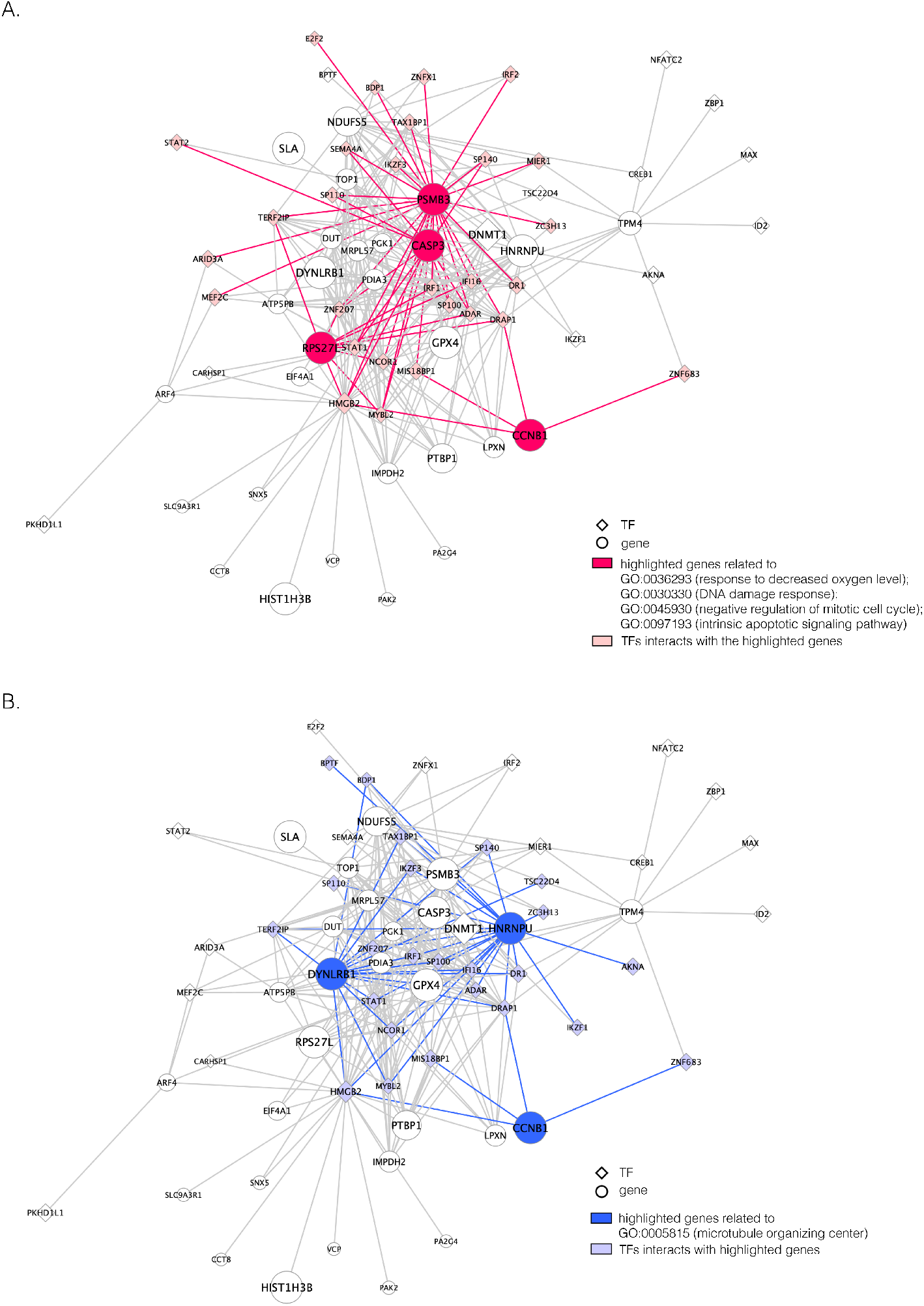
The unique GRNs of DETFs from the patients with severe COVID-19. **A**. GRNs related to: response to a decreased oxygen level (GO:0036293); DNA damage response (GO:0030330); negative regulation of the mitotic cell cycle (GO:0045930); and the intrinsic apoptotic signaling pathway (GO:0097193). **B**. GRNs related to the microtubule organizing center (GO:0005851). The edges are shown if their absolute Pearson correlation coefficients larger than 0.4. DETFs: differentially expressed transcription factors.

## Results

### Effectiveness of neighbor images in removing transitive interactions

We generated simulated data and attempted to train CNNC using the two types of input, one with only the primary images and the other with the augmented images (combined primary and neighbor images, **Methods**). We observed that the overall proportion of false positives and those due to transitive interactions were remarkably decreased by 40.4% and 55.4%, when considering the neighbor images in the model (Fig.1 B). The rationale behind this observation can be regarded as taking a “normalization” on the primary image over their neighborhood to alleviate the overestimation of the strength of interaction. In addition, Fig.1 C and D clearly illustrate that the consideration of neighbor images will not undermine the power in predicting the direct interactions (e.g., gene 1 ⇒ gene 2 in Fig.1 C, and gene 1 ⇒ gene 3 in Fig.1 D). In Fig.1 E, gene 2 connects to gene 3 via the indirect edges gene 2 ⇒ gene 4 ⇒ gene 3. Furthermore, we noticed that the correlations of both {gene 2, gene 4} (|PCC| = 0.81) and gene 4, gene 3 (|PCC| = 0.83) were stronger than the target {gene 2, gene 3} (|PCC| = 0.67), which provided explicit evidence that {gene 2, gene 3} should be marked as a false positive. By considering neighbor images, the model reduce the predicted confidence score of {gene 2, gene 3} from 0.672 to 0.001, with a similar situation observed in Fig.1 F. These findings consolidate the importance of considering the local neighborhood in GRN construction to eliminate false positives due to transitive interactions.

### Overview of DeepDRIM

DeepDRIM is proposed to reconstruct cell-type-specific GRNs from scRNA-seq data with high precision and a low false positive rate. Fig.2 illustrates how DeepDRIM can be used to predict the interaction between gene *a* and gene *b*. First, DeepDRIM converts the joint gene expression of gene *a* and gene *b* into a two-dimensional histogram with 32 by 32 bins (primary image, Fig.2 A), where the intensity of each bin refers to the number of cells falling within it. Second, DeepDRIM constructs 2*n* + 2 neighbor images, where the 2*n* images that refer to the *n* genes have top positive covariance with gene *a* (*a, i*) or gene *b* (*b, j*) and the 2 images represent the self-images (*a, a*) and (*b, b*). These neighbor images are given to the model to capture the neighborhood context of the primary image (Fig.2 B), which provides the key information required to distinguish the direct and transitive interactions. We organize the neighbor images as a tensor rather than an augmented image to achieve better performance on real data (Additional file 1: Figure S2). Third, two CNNs are used to process the primary image (Network A) and the neighbor image tensor (32 by 32 by 2*n*+2) (Network B), respectively (Fig.2 C, **Methods** and Additional file 1: Figure S3). The neural networks are trained by known TF-gene interactions taken from publicly available cell-type-specific ChIP-seq data. Finally, the unknown interactions are predicted by the directed edges with confidence scores (between 0 and 1, Fig.2 D).

### DeepDRIM outperforms the existing algorithms for reconstructing cell-type-specific GRNs

We collected the scRNA-seq datasets from eight cell lines (see **Methods** for the definitions of their abbreviations) and their corresponding ChIP-seq data from two sources [46, 60] to compare DeepDRIM with the existing methods (Table 1) using TF-aware three-fold cross-validation (**Methods**). We first assessed DeepDRIM with PCC, MI, CNNC, and GENIE3; GENIE3 is one of the best algorithms for reconstructing GRNs on scRNA-seq [46] and bulk gene expression data [19, 36].

**TABLE 1.** scRNA-seq datasets from the eight cell lines used in the experiments.

| Cell lines | Genes | Cells | Size of train-<br>ing set | Number of<br>TFs | Pseudo-<br>time |
| --- | --- | --- | --- | --- | --- |
| Bone marrow-derived macrophages [2] | 20,463 | 6,283 | 50,254 | 13 | N |
| Dendritic cells [2] | 20,463 | 4,126 | 28,046 | 16 | N |
| mESC(1) [31] | 24,175 | 2,717 | 154,931 | 38 | N |
| hESC [11] | 17,735 | 758 | 100,720 | 18 | Y |
| mESC(2) [23] | 18,385 | 421 | 94,332 | 18 | Y |
| mHSC(E) [41] | 4,762 | 1,071 | 49,114 | 18 | Y |
| mHSC(GM) [41] | 4,762 | 889 | 43,712 | 18 | Y |
| mHSC(L) [41] | 4,762 | 847 | 48,884 | 18 | Y |

mESC(1): IB10 mouse embryonic stem cells, hESC: human embryonic stem cells, mESC(2): 5G6GR mouse embryonic stem cells , mHSC(E), mHSC(GM) and mHSC(L): three mouse hematopoietic stem cell lines of erythroid lineage, granulocyte-macrophage lineage, and lymphoid lineage.

Our results demonstrate that DeepDRIM outperformed all four methods in the eight cell types, and was significantly better than the second best CNNC (Fig.3 A-D, Additional file 1: Table S1 and S2) with respect to both AUROC (*p*-values ∈ [1.46*E* − 3, 7.63*E* − 6]) and AUPRC (*p*-values [3.42*E* − 3, 7.63*E* − 6]). We also showed that DeepDRIM efficiently eliminated false positives from CNNC in all the eight scRNA-seq datasets (Additional file 1: Figure S4).

To further evaluate the effectiveness of DeepDRIM, we collected six algorithms that have been recently identified with the highest median AUPRC in synthetic networks and Boolean models from BEELINE [46]. Because some of these algorithms require pseudotime-ordered cells, we selected five eligible cell types (**Table 1** and found the six algorithms perform differently for each of them (Additional file 1: Table S3 and S4). We compared the efficiency of DeepDRIM to these algorithms and found that DeepDRIM significantly outperformed all six tested algorithms (Fig.3 E-F). DeepDRIM achieved an average median AUROC of 0.789 and an AUPRC of 0.809 across the five cell types, while the second best methods only achieved an AUROC of 0.591 (Additional file 1: Table S3) and an AUPRC of 0.657 (Additional file 1: Table S4). The TF-specific AUROC and AUPRC are shown in Additional file 2-4.

### DeepDRIM is robust to the quality of scRNA-seq data and the size of the training set

The performance of DeepDRIM can be affected by the quality of scRNA-seq data (the dropout rate and cell number), the number of involved neighbor images, and the size of the training set. To evaluate the robustness of DeepDRIM toward these factors, we first selected the scRNA-seq data from bone marrow-derived macrophages [2] as a template and simulated a series of scRNA-seq data with a range of parameters (**Methods**). Seven scRNA-seq gene expression datasets were generated by subsampling the involved cell numbers (from 20 to 4,000 cells), which in turn changed the resolution of both the primary and neighbor images. We found DeepDRIM to be robust to the low-resolution images when the number of cells was greater than 100 (Fig.4 A). Next, we imputed the dropouts in the template using MAGIC [55] and then randomly masked the entries as dropouts with a range of dropout rates (**Methods**). As shown in Fig.4 B, DeepDRIM demonstrates stable performance in diverse dropout configurations. Third, we compared the performance of DeepDRIM by varying the number of neighbor images input into the model. As a result, we found that the more neighbor images that were involved, the better the performance of DeepDRIM (Fig.4 C). In practice, involving more images would be more computationally costly. In our study, we chose the top 10 genes with the strongest positive covariance with the target TF or gene; thus involving a total of 22 neighbor images (if not specified) to balance the two factors. In addition, to evaluate the effect of the size of the training set, we subsampled 20%, 40%, 60%, 80%, and 100% of the benchmarked TF-gene pairs for training. Our results revealed that the size of the training set did not significantly affect the performance of DeepDRIM (Fig.4 D), and almost reached a plateau when 40% of the training set (including 20,101 TF-gene pairs) was applied.

### Uncovering the variation of B cell-specific GRNs between the patients with mild and severe COVID-19

Patients diagnosed with COVID-19 can have mild or severe acute respiratory distress syndrome, although the underlying molecular mechanisms responsible for these differences remain unknown. We performed a case study to elucidate the differences in B cell-specific GRNs between the patients with mild and severe COVID-19, because the immune responses have been reported to be distinct between the two situations [6]. To this end, we downloaded scRNA-seq data from the bronchoalveolar lavage fluid of six patients with severe symptoms, three patients with mild symptoms, and three healthy controls [34]. The cell type clusters were obtained by SC3[30] and the one belonged to B cells was recognized according to the marker genes provided by the original paper [34]. We extracted validated TF-gene pairs in B cells from the Gene Transcription Regulation Database [58] as the positive pairs, and combined them with the negative pairs from the same TFs and the gene expression from the healthy controls as the training set (**Methods**).

We observed a clear difference in the GRNs between the two types of patients, and also found that the target genes of the DETFs were highly correlated with severe acute respiratory syndrome coronavirus 2 (SARS-CoV-2) infection. First, we observed that DETFs had significantly more targets (*p*-values = 8.50*E* − 4, Wilcoxon rank sum test) in the patients with severe symptoms, suggesting that these DETFs are more “active” in working with their target genes (**Methods** and Fig.5 A-B). Indeed, the DETFs in the patients with severe symptoms had 1.9 times more targets with high confidence (confidence scores ∈ [0.967, 1]; the last bar in Fig.5 A) than the patients with mild symptoms. Next, we focused on the GRNs of DETFs that were unique to the patients with severe symptoms (Fig.5 C, Fig.6 A and B). The informative target genes were selected based on the following two criteria: 1. They should belong to the top 5,000 genes with the highest expression variance in B cells; and 2. they should be ranked in the top 0.1% of the confidence scores of the patients with severe symptoms. The eligible genes were annotated with PageRank scores [21] (**Methods** and Additional file 5) and gene ontology (GO) modules by gene set enrichment analysis (GSEA) [59](**Methods** and Additional file 6). We identified four GO modules that were associated with two common symptoms in patients with COVID-19, hypoxemia and lymphopenia (Fig.5 C, Fig.6 A): 1. response to decreased oxygen levels (GO:0036293; *PMAIP* 1, *CASP* 3, *PSMB*3, *CCNB*1, *p*-values=4.80*E* − 3); 2. DNA damage response (GO:0030330; *PMAIP* 1, *CCNB*1, *RPS*27*L*, *p*-values=1.51*E* − 2); 3. negative regulation of the mitotic cell cycle (GO:0045930; *PSMB*3, *CCNB*1, *RPS*27*L*, *p*-values=1.22*E* − 2); and 4. the intrinsic apoptotic signaling pathway (GO:0097193; *PMAIP* 1, *CASP* 3, *RPS*27*L*, *p*-values=6.29*E* − 3). The patients were reported to have low oxygen levels or hypoxemia without dyspnea [54, 15], both of which were strongly correlated with the GO modules “response to decreased oxygen level” and associated with “the intrinsic apoptotic signaling pathway” [50]. Cao et al. [8] reported that genes related to apoptosis could lead to lymphopenia in patients with COVID-19. Xiong et al. [57] identified differentially expressed genes in peripheral blood mononuclear cells of patients with COVID-19 and healthy controls. These genes were enriched in apoptosis and p53 signaling pathways, both of which could lead to lymphopenia. Among the genes in these four GO modules, *PMAIP* 1 [28, 47], *CASP* 3 [16, 35], *PSMB*3 [56], and *CCNB*1 [57] have been reported to be associated with COVID-19 individually (Additional file 1: Table S5).

In addition to these main findings, we also noted that there were four genes with top PageRank scores in the patients with severe symptoms in which unique GRNs could be related to SARS-CoV-2 infection. Three of them (*DY NLRB*1, *HNRNPU*, and *CCNB*1) belong to GO:0005815 (microtubule organizing center, *p*-values=5.33*E* − 3), which has been reported to be a major facilitator of virus infection [18] due to its ability to provide invading pathogens with directed transport (Fig.6 B). The other gene *DNMT* 1 is related to *ACE*2 [49], which is a known co-receptor for the SARS-CoV-2 [42].

## Discussion

Understanding the GRNs is fundamental to the advancement of molecular biology research. Gene expression profiles from high-throughput sequencing enable computational algorithms to reconstruct GRNs by examining TF-gene co-expression. Bulk RNA-seq hides the gene activities at single-cell resolution and will be replaced by scRNA-seq in the near future. However, the gene expression distribution from scRNA-seq data is not consistent with the assumptions made by most of the existing methods, which leads to their poor performance in reconstructing GRNs on the scRNA-seq data [33]. In addition, the widely spread dropouts cause bias in calculating gene-gene co-expression, even after imputation [12].

In this study, we propose DeepDRIM, a supervised deep neural network, to reconstruct GRNs on scRNA-seq data. Comprehensive evaluation of the performance of DeepDRIM on different cell types demonstrated that it outperformed the existing algorithms designed for either bulk or scRNA-seq gene expression data. It is inadvisable to calculate TF-gene interactions on scRNA-seq data using classical correlation-based methods due to the ubiquitous cellular heterogeneity and dropouts (Fig.3 A-D). To avoid these limitations, DeepDRIM converts the numerical representation of TF-gene expression to an image and applies a CNN to embed it into a lower dimension. This strategy also avoids data normalization and does not presume any distribution. DeepDRIM requires validated TF-gene pairs for use as a training set to highlight the key areas in the embedding space that can distinguish the direct interactions and false positives.

We trained and tested DeepDRIM using data from the same cell type. As there is sometimes an insufficient number of cells or validated TF-gene pairs in the training set, we were interested in training the model using one cell type and then applying it to another. We trained DeepDRIM using bone marrow-derived macrophages and then applied it to mESC(1) and vice versa (Additional file 1: Figure S5). The results suggest that it is necessary to apply DeepDRIM to matched cell types in training and test sets; thus, ideas such as transfer learning between cell types are not applicable to this supervised model.

The neighborhood context of the target TF-gene pairs has been widely applied to remove false positives in GRN reconstruction from bulk gene expression data via z-score normalization [17], conditional MI [61, 62], and graphical lasso [13]. However, these methods commonly assume that the gene expression profiles follow a Gaussian distribution, which violates our observation in scRNA-seq data. Most of the existing algorithms designed for scRNA-seq are unsupervised and require pseudotime-ordered cells, making them inapplicable to bone marrow-derived macrophages, dendritic cells, and mESC(1), as illustrated in Table 1. DeepDRIM uses the neighborhood context with respect to neighbor images, and consists of two parts: 1) images from the genes that positively correlate with the TF or gene from the target pair, and 2) two self-images. In the current model, we adopted covariance to select the top correlated genes. Although such linear correlation is not resistant to outliers and dropouts, similar method has shown its effectiveness in discovering gene-gene co-expression from scRNA-seq data [5]. The two self-images can highlight the variance of single gene expression.

DeepDRIM can not only predict the existence of TF-gene interactions, but also determine their causalities. This task is not given much attention by the unsupervised algorithms, despite it being an important consideration if regulatory interactions exist between two TFs. For this particular task, DeepDRIM does not surpass CNNC, because CNNC only focuses on the primary image and it is easier to capture the causalities by learning the regulatory directions from the validated TF-gene pairs. We generated a combined model from DeepDRIM and CNNC (Additional file 1: Supplementary Notes) and found that it can effectively reduce the false positives without losing any accuracy in the prediction of causality (Additional file 1: Figure S6).

Many studies have been proposed with the aim to identify all of the cell types in the human tissues, with the ultimate goal of creating a human cell atlas to facilitate interpretation of the gene activities in individual cell types. DeepDRIM bridges the gap between cell types and gene functions, and will serve to increase our understanding of the activities of key TFs. We believe that as the cell-type-specific ChIP-seq data accumulate, DeepDRIM will attract increased attention in the scRNA-seq research community, and will shed light on drug target discovery and precision medicine in the future.

## Conclusion

We propose DeepDRIM, a supervised deep neural network model, to predict GRNs from scRNA-seq data. DeepDRIM converts the joint expression of a TF–gene pair into a primary image and considers the neighbor images as the neighborhood context of the primary image to remove false positives due to transitive interactions. DeepDRIM also utilizes the training set to capture the key areas in the CNN embeddings that can recognize the TF-gene interactions and causalities. Our findings demonstrate that DeepDRIM outperforms nine existing algorithms on the eight cell types tested and is robust to the quality of scRNA-seq data. DeepDRIM can also identify the GRNs of B cells that are different between patients with mild and severe COVID-19 symptoms. We believe that DeepDRIM can fill the gaps in reconstructing cell-type-specific GRNs on scRNA-seq data and contributes to the rapidly growing single-cell research community.

## Methods

### Representation of gene pair joint expression

The scRNA-seq gene expression profiles are represented as a two-dimensional matrix *M*, where *M*_*g,c*_ represents the expression of gene *g* in cell *c*. We added a small pseudo-count to *M*_*g,c*_ to avoid empty entries before applying log-normalization:

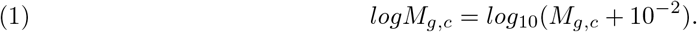

The joint histogram of genes *i* and *j* (*H*_*i,j*_) is generated by splitting *logM*_*i,−*_ and *logM*_*j,−*_ (“-”: across all of the cells) into 32 bins, respectively. The value of each bin is derived from the number of cells that falls in the corresponding slot; this value is further log-normalized to avoid extreme values:

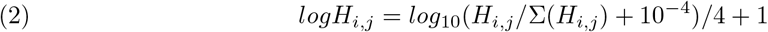

We generated an image (*I*_*i,j*_) for genes *i* and *j* of 32 by 32 pixels, where the intensity of each pixel is the corresponding value in *logH*_*i,j*_. DeepDRIM requires two image sets to predict the direct interaction between genes *i* and *j*, namely 1. the primary image *I*_*i,j*_ and 2. the neighbor images. The neighbor images consist of 1. 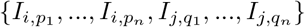, where (*p*_1_*, p*_2_*, ….p*_*n*_) and (*q*_1_*, q*_2_*,…q*_*n*_) are the top *n* genes that have strong positive covariance with gene *i* and gene *j*, respectively; and 2. two self-images *I*_*i,i*_ and *I*_*j,j*_. The default value of *n* was 10 in the experiments.

### Network structure of DeepDRIM

The network structure of DeepDRIM consists of two components, Network A and Network B, which process the primary and neighbor images, respectively (Fig.2 C and Additional file 1: Figure S3). Network A is inspired by VGGnet [52], which contains the stacked convolutional and maxpooling layers, and uses the rectified linear activation function (*ReLu*) as the activation function. The structure of Network B is similar to that of Network A, and is a siamese-like neural network, where the weights are shared among all of the subnetworks. Each image is embedded into a vector of size 512, and a total of 2*n* + 3 images (1 primary image and 2*n* + 2 neighbor images) are converted into a vector of size 512 × (2*n* + 3). This vector is then condensed by two stacked fully connected layers, and is processed for binary classification using the sigmoid function. Moreover, DeepDRIM is trained by mini-batched stochastic gradient descent.

### Simulation of scRNA-seq data to examine the effect of neighbor images

We simulated 2,500 small datasets, each with 4 genes and 1,000 cells. The ground truth network for each dataset was represented by a sparse precision matrix Θ, where each entry had a 50% chance of being non-zero and drawn from [−1, −0.25] ⋃ [0.25, 1], or otherwise was assigned zero. We simulated the gene expression profiles from a multivariate normal distribution *N* (0, Θ^*−*1^) [14]. Next, we randomly chose two gene pairs from each dataset, one involving a direct interaction (Θ_*i,j*_ ≠ 0) as a positive case, and the other involving an independent pair (Θ_*i,j*_ = 0) as a negative case. For each case, we prepared two types of images, a primary image of 32 by 32 pixels, and an augmented image by concatenating the primary and six neighbor images (Fig.1 C–F) of 96 by 96 pixels. We generated two training sets with 5,000 primary and 5,000 augmented images, respectively. These images were used to train CNNC and the performance was evaluated using the AUROC from the five-fold cross-validation.

### scRNA-seq data from eight cell lines

We prepared the real scRNA-seq data from eight cell lines and the corresponding cell-type-specific ChIP-seq data as the benchmarks (Table 1) to compare DeepDRIM with the existing algorithms for GRN reconstruction. The eight cell lines comprised bone marrow-derived macrophages [2], dendritic cells [2], IB10 mouse embryonic stem cells (mESC(1)) [31], human embryonic stem cells (hESC) [11], and 5G6GR mouse embryonic stem cells (mESC(2)) [23], as well as three mouse hematopoietic stem cell lines [41] of erythroid lineage (mHSC(E)), granulocyte-macrophage lineage (mHSC(GM)), and lymphoid lineage (mHSC(L)). All scRNA-seq data were pre-processed and normalized according to the descriptions in [46, 60]. In practice, GENIE3 is slow if too many genes or cells are involved; thus, we removed the less informative cells and genes using the strategies described in [1].

We extracted the validated TF targets from the ChIP-seq data as positive cases, and the same number of non-validated targets as negative cases. As training sets that are too large and are computationally insolvable in terms of generating images, we randomly selected 18 TFs and their validated targets as positive cases in the training data for hESC, mESC(2), mHSC(E), mHSC(GM), and mHSC(L) to alleviate the computational burden (Table 1).

To improve the performance of the unsupervised methods in Fig.3 E-F, only the overlap between top-varying 500 genes and the TFs/genes in the training set were selected from the scRNA-seq data of hESC, mESC(2), mHSC(E), mHSC(GM) and mHSC(L). In cross-validation, We trained CNNC and DeepDRIM using 2/3 TF-gene pairs in the training set and evaluated their performance on the overlap between the TFs/genes in the remaining 1/3 test set and top-varying 500 genes. This could guarantee all the supervised and unsupervised were evaluated on the same TF-gene pairs.

### Comparison of DeepDRIM to existing algorithms for GRN reconstruction

We compared DeepDRIM with the nine existing algorithms using their default parameters. The nine algorithms were PCC, MI, CNNC [60], PIDC [9], GENIE3[26], GRNBOOST2 [39], SCODE [38], PPCOR [29], and SINCERITIES [44]. With the exception of PCC, MI, and CNNC, the other six methods were performed using the interfaces provided by BEELINE [46]. The AUROC and AUPRC for each TF were collected to calculate the *p*-values between two algorithms using the Wilcoxon signed rank test. Given that CNNC and DeepDRIM are supervised models, the TFs from the ChIP-seq data were divided into three independent parts for cross-validation (Additional file 1: Supplementary Note).

### Simulation of scRNA-seq data to evaluate robustness

The simulated datasets were transferred from the scRNA-seq of bone marrow-derived macrophages [2] to preserve the characteristics of scRNA-seq data. We simulated gene expression profiles with various cell numbers and sizes of training sets via sub-sampling from the total 6,283 cells and 50,254 validated TF-gene pairs from the ChIP-seq data. We applied MAGIC [55] to impute the missing values in the raw gene expression matrix, and subsequently masked the corresponding entries according to the “dropout step” in BoolODE [46]. BoolODE has two parameters, *drop − probability* and *drop − cutoff*, which are used to control the number of entries to be masked. The entries have a probability of *drop − probability* to be masked if their gene expression values are at the bottom *drop − cutoff*. We set the *drop − probability* = 0.3, 0.5 and the *drop − cutoff* = 0 to 0.9.

### Generation of validated TF-gene pairs for B cells in patients with COVID-19

We extracted the ChIP-seq data with the keyword “human B cell” in the Gene Transcription Regulation Database [58] and determined the TF target genes as those with high confidence peaks (*p*-value *<* 1*E* − 8) in the promoter regions of these genes. The promoter regions were defined as the 10 kb upstream and 1 kb downstream regions of the transcript start sites. To generate a balanced training set, we extracted an equal number of negative pairs by randomly selecting the non-target genes of the selected TFs.

### Identification of differentially expressed TFs

We applied SCDE [27] to determine the differentially expressed TFs if the expression fold changes *>* 2 or *<* 0.5, and the *p*-values to be *<* 1*E −* 11 after multiple testing correction.

### Gene PageRank score and functional annotation

We calculated gene PageRank scores using “networkx” [21] (Additional file 5) and applied GSEA to annotate the enriched GO modules with *p*-value*<* 0.05[59]. The genes were ordered by their PageRank scores in GSEA analysis.

## Availability of data and materials

DeepDRIM is available at https://github.com/jiaxchen2-c/DeepDRIM. Gene expression and ChIP-Seq data of bone marrow-derived macrophages, dendritic cells, mESC(1) are available at https://github.com/xiaoyeye Gene expression and ChIP-Seq data of hESC, mESC(2), mHSC(E), mHSC(GM), mHSC(L) are available at https://doi.org/10.5281/zenodo.3378975. Gene expression profiles from the bronchoalveolar lavage fluid of COVID-19 patients and healthy controls are available at GSE145926.

## Authors’ contributions

LZ, WKC conceived the study; LZ, JXC designed DeepDRIM; JXC implemented the algorithm and analyzed the results. JXC, CWC conducted the experiments. JXC, LZ, WKC wrote the article. JML, APL and ZX reviewed the paper. All authors read and approved the final manuscript.

## Acknowledgements

We also thank Research Grants Council of Hong Kong, Hong Kong Baptist University and HKBU Research Committee for their kind support of this project.

## Additional Files

Additional file 1 — Supplementary Note, Supplementary Figures and Supplementary Tables.

Additional file 2 — AUROCs and AUPRCs of PCC, MI, GENIE3, CNNC, and DeepDRIM for each TF on bone marrow-derived macrophages, dendritic cells, and mESC(1).

Additional file 3 — AUROCs and AUPRCs of PCC, MI, GENIE3, CNNC, and DeepDRIM for each TF on hESC, mESC(2), mHSC(E), mHSC(GM), and mHSC(L).

Additional file 4 — AUROCs and AUPRCs of PIDC, GENIE3, GRNBOOST2, SCODE, PPCOR, SINCERITIES, and DeepDRIM for each TF on hESC, mESC(2), mHSC(E), mHSC(GM), and mHSC(L).

Additional file 5 — PageRank scores, degree and betweenness of the genes in the GRNs from the patients with severe COVID-19.

Additional file 6 — GO annotation for the genes in the GRNS from the patients with severe COVID-19.

