## Additional_file1 for "DeepDRIM: a deep neural network to reconstruct cell-type-specific gene regulatory network using single-cell RNA-Seq Data"

### Supplementary Note

#### Evaluate CNNC and DeepDRIM by three-fold cross-validation

Because CNNC and DeepDRIM are supervised models, we adopted three-fold cross-validation to assess their performance. We kept a balanced positive and negative pairs for each TF and divided these TFs into three partitions, each kept comparable total TF-gene pairs. For example, in bone marrow-derived macrophages, we had total 13 TFs from ChIP-seq data and divided them into three partitions with 4,4 and 5 TFs, respectively. We would carefully adjusted the assignment of TFs to make the numbers of TF-gene pairs in the three partitions comparable.

#### Combined model for causality prediction

We tried to generate a combined model to effectively remove the transitive interactions and infer their causalities simultaneously. We changed the final prediction layer with ”softmax” function, and made label ”0” to represent no interaction between genes $g_a$ and $g_b$, label ”1” to represent $g_a$ regulates $g_b$, and label 2 represent $g_b$ regulates $g_a$. We divided the prediction into two subtasks: 1. whether the label is ”0” (TF-gene has interaction or not); 2. whether the label is ”1” or ”2” (infer the causality). The combined model uses DeepDRIM to deal with the subtask 1 and adopts CNNC to predict the causality for the subtask 2.

#### Supplementary Figures

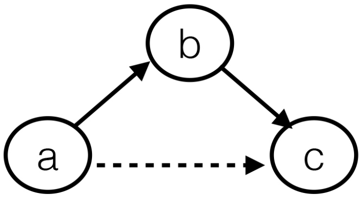


**Figure S1** An example of transitive interaction. Gene **a** and gene **c** strongly correlate with each other through an intermediate gene **b**.

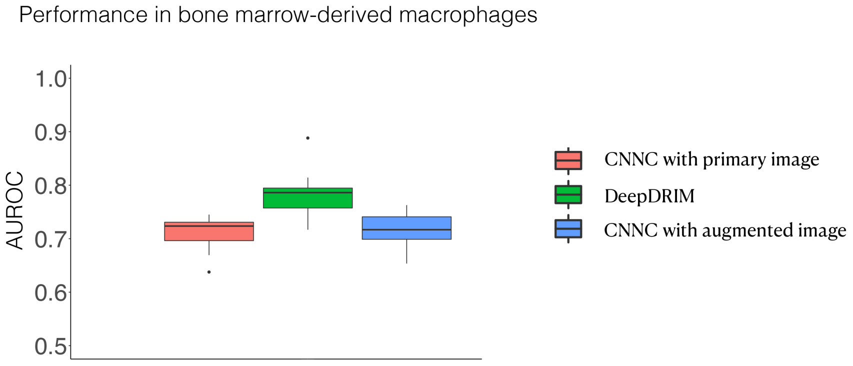


**Figure S2** The performance of CNNC with primary image and augmented image as inputs and DeepDRIM.

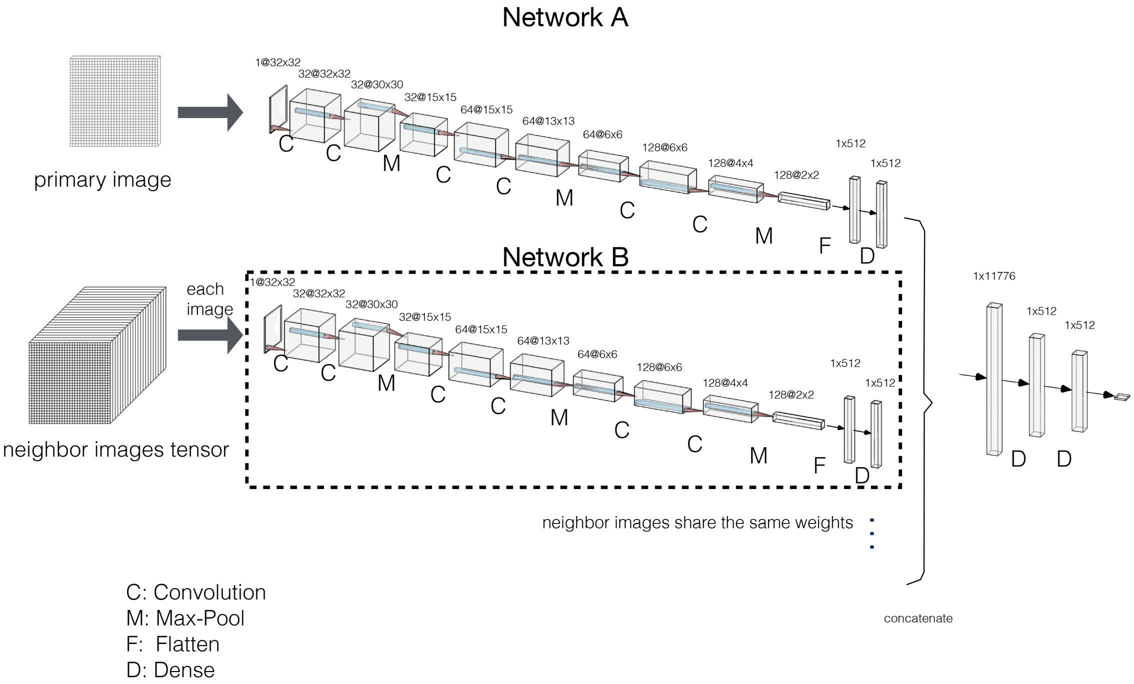


**Figure S3** Network structure of DeepDRIM.

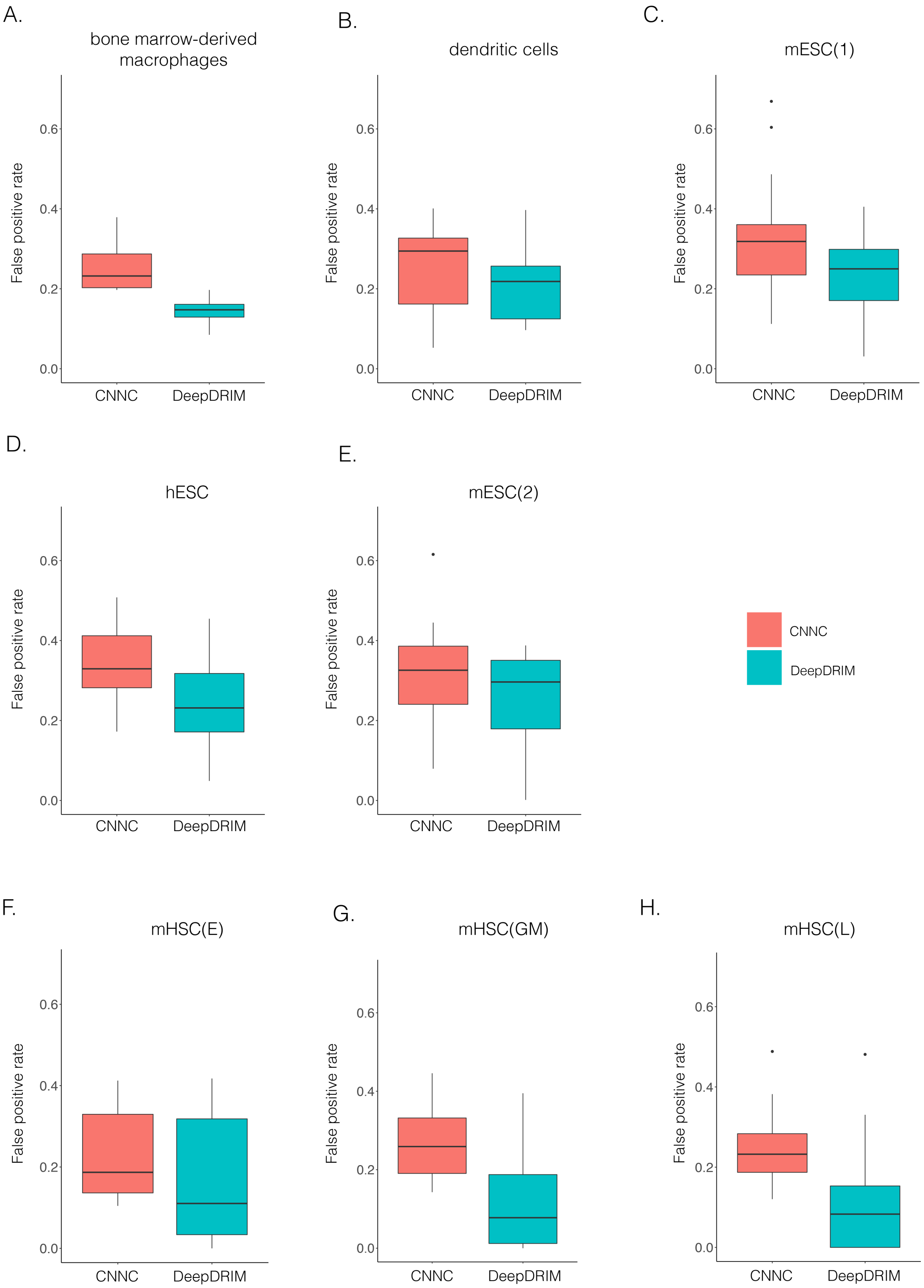


**Figure S4** False positive rates of CNNC and DeepDRIM for the eight cell types. The false positive rates are calculated by considering the interactions whose confidence scores are in top 10% of the corresponding algorithms. We excluded the TFs with less than 20 targets for better capturing of false positive rates.

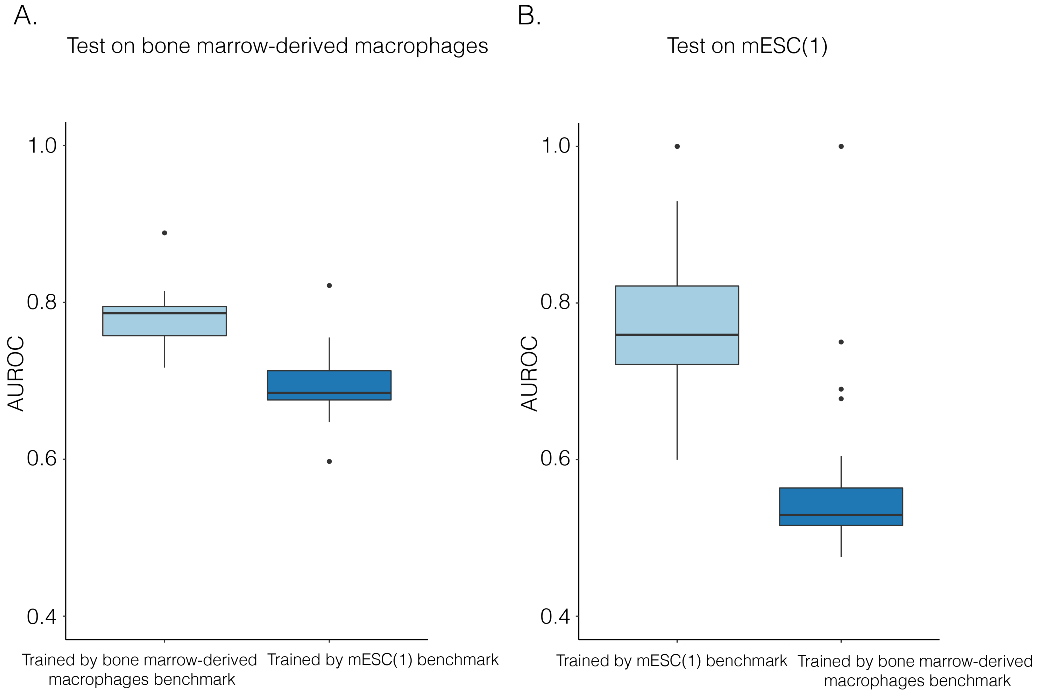


**Figure S5** The performance of DeepDRIM that trained and tested on the same and different cell types.

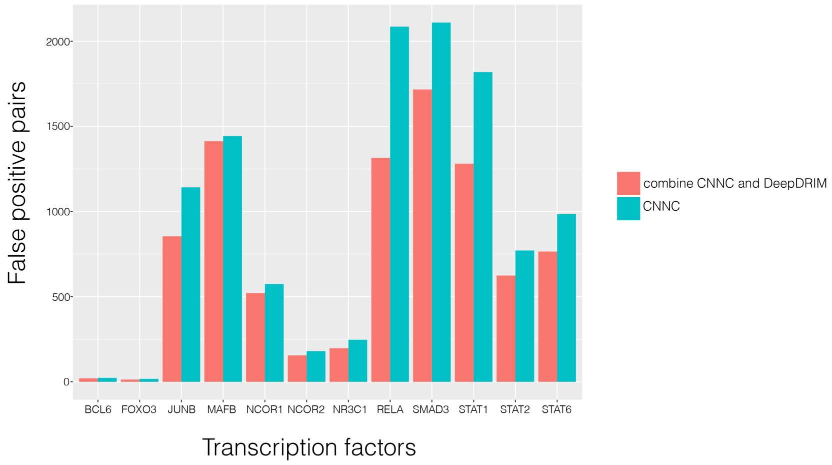


**Figure S6** The performance of CNNC and the combined model on bone marrow-derived macrophages by considering both the existences and causalities of TF-gene interactions.

#### Supplementary Tables

**Table S1** Median AUROC of the involved TFs for PCC, MI, GENIE3, CNNC, and DeepDRIM. The algorithms' ranks are shown in the parentheses.

| | bone marrow-derived macrophages | mESC(1) | dendritic cells | hESC | mESC(2) | mHSC(E) | mHSC(GM) | mHSC(L) |
|---|---|---|---|---|---|---|---|---|
| PCC | 0.628(4) | 0.465(5) | 0.626(4) | 0.542(3) | 0.538(4) | 0.477(5) | 0.561(4) | 0.546(5) |
| MI | 0.684(3) | 0.642(3) | 0.734(3) | 0.539(4) | 0.563(3) | 0.575(3) | 0.600(3) | 0.592(3) |
| GENIE3 | 0.584(5) | 0.495(4) | 0.582(5) | 0.516(5) | 0.477(5) | 0.532(4) | 0.557(5) | 0.558(4) |
| CNNC | 0.724(2) | 0.695(2) | 0.736(2) | 0.638(2) | 0.676(2) | 0.730(2) | 0.724(2) | 0.747(2) |
| DeepDRIM | **0.786(1)** | **0.756(1)** | **0.748(1)** | **0.762(1)** | **0.724(1)** | **00.804(1)** | **0.834(1)** | **0.873(1)** |

**Table S2** Median AUPRC of the involved TFs for PCC, MI, GENIE3, CNNC and DeepDRIM. The algorithms' ranks are shown in the parentheses.

| | bone marrow-derived macrophages | mESC(1) | dendritic | hESC | mESC(2) | mHSC(E) | mHSC(GM) | mHSC(L) |
|---|---|---|---|---|---|---|---|---|
| PCC | 0.644(4) | 0.509(5) | 0.638(4) | 0.543(3) | 0.562(4) | 0.494(5) | 0.540(4) | 0.545(4) |
| MI | 0.672(3) | 0.613(4) | 0.693(2) | 0.534(4) | 0.573(3) | 0.550(3) | 0.576(3) | 0.584(3) |
| GENIE3 | 0.611(5) | 0.615(3) | 0.615(5) | 0.528(5) | 0.557(5) | 0.511(4) | 0.518(5) | 0.525(5) |
| CNNC | 0.700(2) | 0.658(2) | 0.686(3) | 0.631(2) | 0.625(2) | 0.694(2) | 0.665(2) | 0.708(2) |
| DeepDRIM | **0.775(1)** | **0.710(1)** | **0.709(1)** | **0.726(1)** | **0.661(1)** | **0.769(1)** | **0.802(1)** | **0.851(1)** |

**Table S3** Median AUROC of the involved TFs for PIDC, GENIE3, GRNBOOST2, SCODE, PPCOR, SINCERITIES, and DeepDRIM. The algorithms' ranks are shown in the parentheses.

| | hESC | mESC(2) | mHSC(E) | mHSC(GM) | mHSC(L) |
|---|---|---|---|---|---|
| PIDC | 0.499(7) | 0.641(2) | 0.468(6) | 0.490(7) | 0.502(5) |
| GENIE3 | 0.626(2) | 0.341(7) | 0.547(2) | 0.597(2) | 0.544(2) |
| GRNBOOST2 | 0.615(3) | 0.399(6) | 0.529(3) | 0.527(3) | 0.534(3) |
| SCODE | 0.500(5) | 0.459(4) | 0.506(5) | 0.515(4) | 0.477(7) |
| PPCOR | 0.500(5) | 0.495(3) | - | 0.499(5) | 0.5(6) |
| SINCERITIES | 0.538(4) | 0.457(5) | 0.507(4) | 0.495(6) | 0.520(4) |
| DeepDRIM | **0.704(1)** | **0.875(1)** | **0.755(1)** | **0.793(1)** | **0.818(1)** |

PPCOR failed to run on mHSC(E) due to an unexpected matrix singularity error.

**Table S4** Median AUPRC of the involved TFs for PIDC, GENIE3, GRNBOOST2, SCODE, PPCOR, SINCERITIES, and DeepDRIM. The algorithms' ranks are shown in the parentheses.

| | hESC | mESC(2) | mHSC(E) | mHSC(GM) | mHSC(L) |
|---|---|---|---|---|---|
| PIDC | 0.599(5) | 0.609(2) | 0.604(6) | 0.609(5) | 0.644(4) |
| GENIE3 | 0.679(2) | 0.376(7) | 0.684(2) | 0.651(2) | 0.639(5) |
| GRNBOOST2 | 0.666(3) | 0.436(6) | 0.631(4) | 0.631(3) | 0.657(3) |
| SCODE | 0.581(7) | 0.444(5) | 0.632(3) | 0.631(3) | 0.615(7) |
| PPCOR | 0.589(6) | 0.481(3) | - | 0.601(7) | 0.632(6) |
| SINCERITIES | 0.638(4) | 0.471(4) | 0.624(5) | 0.606(6) | 0.664(2) |
| DeepDRIM | **0.750(1)** | **0.810(1)** | **0.800(1)** | **0.829(1)** | **0.856(1)** |

PPCOR failed to run on mHSC(E) due to an unexpected matrix singularity error.

**Table S5** Descriptions for the genes with top PageRank scores in the unique GRNs from the patients with severe COVID-19. ($Y$ denote direct evidence and $I$ denote indirect evidence)

| gene symbol | rank of PageRank value | keyword | description | enriched GO modules | associated with COVID-19 | citations |
|---|---|---|---|---|---|---|
| PMAIP | 1 | apoptosis | $PMAIP1$ has also been recently found to be related to COVID-19 [1, 2] by Apoptosis. According to their studies, $PMAIP1$ promotes proteasomal degradation of $MCL1$, where $MCL1$ and $PMAIP1$ are found significantly altered after SARS-CoV or HCoV-229E infection. | GO:0036293 (response to decreased oxygen level, $p$-values=$4.05E-3$). GO:0030330 (DNA damage response, $p$-values=$1.26E-2$). GO:0097193 intrinsic apoptotic signaling pathway, $p$-values=$6.01E-3$) | Y (yes) | [1, 2] |
| CASP3 | 1 | apoptosis | $CASP3$ play an important role in the execution-phase of cell apoptosis. $CASP3$ is also founded as one of apoptosis-related genes that have increase expression in scRNA profiles in M1 phenotype macrophages in the coculture model to study interaction among macrophages, lung cells and SARS-CoV-2 [3]. $CASP3$ is also listed as a key target in the drug-disease common targets when exploring the pharmacology about COVID-19 [4]. | GO:0036293 (response to decreased oxygen level, $p$-values=$4.05E-3$). GO:0097193 (intrinsic apoptotic signaling pathway, $p$-values=$6.01E-3$) | Y | [3, 4] |
| PIM3 | 1 | apoptosis | $PIM3$ is related to the pathway of Apoptosis and Autophagy, and can regulate $AMPK$'s activities, while $AMPK$ may decrease $ACE$ expression [5]. $ACE$'s novel homolog angiotensin converting enzyme 2 (ACE2) is known as the co-receptor for the coronavirus and plays an important role in SARS-CoV-2 infection [6]. | GO:0007346 (regulation of mitotic cell cycle, $p$-values=$4.67E-4$) | I (indirect association) | [5, 6] |
| GPX4 | 1 | T cell | $GPX4$ can protect T cell from ferroptosis and support T cell expansion, thus are associated with primary T cell response to viral and parasitic infection | GO:0055114 (oxidation-reduction process, $p$-values=$7.36E-4$) | - | - |
| DYNLB1 | 1 | microtubule | $DYNLB1$ is a member of the roadblock dynein light chain family. The encoded cytoplasmic protein is capable of binding intermediate chain proteins, interacts with transforming growth factor-beta, and has been implicated in the regulation of actin modulating. Several viruses are known to interact with tubulin or their molecular motors like kinesin or dynein proteins. | GO:0005815 (microtubule organizing center, $p$-values=$4.05E-3$) | - | - |
| PSMB3 | 1 | remove damaged proteins | $PSMB3$ is a protein coding gene which is related to removing misfolded or damaged proteins, and is identified as a gene in the gene set for proteotoxic stress which suggest the terminal exhaustion [7], and the gene set is found dominated in critical COVID-19 compare to mild, implying the inflammation-driven terminal exhaustion and severe dysregulation. | GO:0036293 (response to decreased oxygen level, $p$-values=$4.05E-3$). GO:0045930 (negative regulation of mitotic cell cycle, $p$-values=$1.08E-2$) | Y | [7] |

| | | | | | | |
|---|---|---|---|---|---|---|
| DNMT1 | 2 | ACE2 | *DNMT1* is related with *ACE2*, thus affects SARS-CoV-2 infection through DNA methylation and chromatin silencing [8]. | GO:0010638 (positive regulation of organelle organization, *p*-values=$1.05E-3$) | I | [8] |
| SLA | 3 | T cell | *SLA* negatively regulates T cell receptor (TCR) signaling, where TCR has recently found to be correlated with COVID-19 [9]. | GO:0050896 (response to stimulus, *p*-values=$6.51E-3$) | I | [9] |
| HNRNPU | 4 | microtubule | *HNRNPU* are involved in the formation of stable mitotic spindle microtubules (MTs) attachment to kinetochore, spindle organization and chromosome congression [10]. | GO:0005815 (microtubule organizing center, *p*-values=$4.05E-3$) | - | - |
| CCNB1 | 5 | apoptosis and P53 signaling, microtubule | *CCNB1*, is a protein coding gene and are involved in mitosis as well as maturation-promoting factor (MPF), is a necessary for the control of G2/M transition phase in cell cycle, and is identified as one of the significantly altered genes that enriched to the apoptosis and P53 signaling [11], which may be related to the reducing of Lymphocytes in COVID-19 patients. | GO:0030330 (DNA damage response, *p*-values=$1.26E-2$). GO:0045930 (negative regulation of mitotic cell cycle, *p*-values=$1.08E-2$). GO:0005815 (microtubule organizing center, *p*-values=$4.05E-3$) | Y | [11] |
| RPS27L | 5 | cell apoptosis | *RPS27L* is related to cysteine-type endopeptidase activator activity involved in apoptotic process. | GO:0030330 (DNA damage response, *p*-values=$1.26E-2$). GO:0045930 (negative regulation of mitotic cell cycle, *p*-values=$1.08E-2$)GO:0097193 (intrinsic apoptotic signaling pathway, *p*-values=$6.01E-3$) | - | - |
| HIST1H3B (H3C2) | 5 | histones | H3C2 is a Protein Coding gene related to Histones, while Histones are basic nuclear proteins responsible for the nucleosome structure. Therefore it is important in transcription regulation, DNA repair, DNA replication and chromosomal stability. | GO:0010608 (posttranscriptional regulation of gene expression, *p*-values=$2.63E-3$) | - | - |

[2] Shitao Rao, Alexandria Lau, and Hon-Cheong So. Exploring diseases/traits and blood proteins causally related to expression of ace2, the putative receptor of sars-cov-2: A mendelian randomization analysis highlights tentative relevance of diabetes-related traits. *Diabetes Care*, 2020.

[3] Fuyu Duan, Liyan Guo, Liuliu Yang, Yuling Han, Abhimanyu Thakur, Benjamin E Nilsson-Payant, Pengfei Wang, Zhao Zhang, Chui Yan Ma, Xiaoya Zhou, et al. Modeling covid-19 with human pluripotent stem cell-derived cells reveals synergistic effects of anti-inflammatory macrophages with ace2 inhibition against sars-cov-2. 2020.

[4] Xiao-Ying Ling, Jia-Lei Tao, Xun Sun, and Bin Yuan. Exploring material basis and mechanism of lianhua qingwen prescription against coronavirus based on network pharmacology. *Chin. Trad. Herbal Drugs*, pages 1723–1730, 2020.

[5] Jia Liu, Xuan Li, Qingguo Lu, Di Ren, Xiaodong Sun, Thomas Rousselle, Ji Li, and Jiyan Leng. Ampk: a balancer of the renin–angiotensin system. *Bioscience Reports*, 39(9):BSR20181994, 2019.

[6] Andrew M South, Debra I Diz, and Mark C Chappell. Covid-19, ace2, and the cardiovascular consequences. *American Journal of Physiology-Heart and Circulatory Physiology*, 2020.

[7] Els Wauters, Pierre Van Mol, Abhishek D Garg, Sander Jansen, Yannick Van Herck, Lore Vanderbeke, Ayse Bassez, Bram Boeckx, Bert Malengier-Devlies, Anna Timmerman, et al. Discriminating mild from critical covid-19 by innate and adaptive immune single-cell profiling of bronchoalveolar lavages. *BioRxiv*, 2020.

[8] Amr H Sawalha, Ming Zhao, Patrick Coit, and Qianjin Lu. Epigenetic dysregulation of ace2 and interferon-regulated genes might suggest increased covid-19 susceptibility and severity in lupus patients. *Clinical Immunology*, page 108410, 2020.

[9] Lucas Gutierrez, John Beckford, and Houda Alachkar. Deciphering the tcr repertoire to solve the covid-19 mystery. *Trends in Pharmacological Sciences*, 2020.

[10] Nan Ma, Sachihiro Matsunaga, Akihiro Morimoto, Gyosuke Sakashita, Takeshi Urano, Susumu Uchiyama, and Kiichi Fukui. The nuclear scaffold protein saf-a is required for kinetochore–microtubule attachment and contributes to the targeting of aurora-a to mitotic spindles. *Journal of cell science*, 124(3):394–404, 2011.

[11] Yong Xiong, Yuan Liu, Liu Cao, Dehe Wang, Ming Guo, Ao Jiang, Dong Guo, Wenjia Hu, Jiayi Yang, Zhidong Tang, et al. Transcriptomic characteristics of bronchoalveolar lavage fluid and peripheral blood mononuclear cells in covid-19 patients. *Emerging microbes & infections*, 9(1):761–770, 2020.
